# Task-specific ISS reduction and network balance during Stroop task performance in older adults

**DOI:** 10.64898/2026.07.13.738225

**Authors:** Kazuya Ouchi, Haruki Yokota, Narihisa Matsumoto, Tomokazu Tsurugizawa

## Abstract

Declines in the cognitive inhibition that comes with aging impacts daily life and independence. It is still debated whether overactivation during cognitive inhibition tasks in older adults is due to compensation or neural noise. To address this question, we examined age-related changes in inter-subject similarity (ISS) of functional connectivity from task-based and resting-state fMRI. Using Bayesian hierarchical modeling, we identified 27 Stroop task-related regions of interest and found that ISS in these regions was significantly reduced in older adults during task performance. Furthermore, aging is associated with a loss of consistent functional connectivity patterns in frontal regions, accompanied by the emergence of a convergent compensatory mechanism within visual attention regions. Principal component analysis of individual activation deviations from the group-mean pattern showed that task performance in older adults cannot be explained by overall task-related brain activation, but rather by a specific spatial component involving suppression of the default mode network and increased activity in visual attention regions. These findings indicate that task-related brain overactivation in older adults is not due to uniform noise or uniform compensation, but rather a spatially specific pattern of functional reorganization.

## 1. Introduction

Cognitive inhibition is an important aspect of executive function, which is a part of higher cognitive function (*1, 2*). Cognitive inhibition plays a role in supporting appropriate behavioral selection in daily life (*3, 4*), and its decline in older adults leads to a decreased quality of life, including an increased risk of falls (*5, 6*). Therefore, understanding age-related decline in cognitive inhibition is required in an aging society.

The Stroop task is a representative paradigm for assessing cognitive inhibition (*7*), in which participants were required to name the ink color of the presented word, rather than reading the word’s meaning. In the incongruent condition, in which the word meaning and ink color did not match, the response time is prolonged compared to that in the congruent condition. This phenomenon is more pronounced in older than younger adults (*8*). Functional magnetic resonance imaging (fMRI) studies have consistently reported activation of the anterior cingulate cortex, dorsolateral prefrontal cortex, and parietal cortex during the Stroop task in younger adults (*9*). Furthermore, in older adults, increased activation has been observed not only in these regions but also in additional brain regions, such as the inferior frontal gyrus and supplementary motor area, compared to young adults (*10, 11*). However, the cause of this overactivation is still debated. Some research has indicated a “compensation” for functional decline to maintain cognitive performance (*12*). Another study suggests the “noise” attributable to reduced neural efficiency (*13*). Distinguishing these interpretations requires examining inter-individual consistency in overactivation.

Inter-subject similarity (ISS) is a useful index for examining the stability of functional connectivity (FC) patterns across individuals. A high ISS indicates common neural mechanisms across individuals, whereas a low ISS indicates variable processing strategies. A previous study with resting state and task fMRI showed that older adults have a significantly lower ISS than younger adults (*14*). However, most studies have used the ISS analyses at the whole-brain level, which inevitably includes signals from regions unrelated to the task. Therefore, it is essential to identify task-related regions and then perform ISS analyses within those regions. A general linear model (GLM) is conventionally used to identify regions activated during task in fMRI. However, this method has a difficulty detecting regions with large individual differences in activation patterns. To overcome this limitation, we employed hierarchical Bayesian modeling to estimate activation because this model regards individual parameters as deviations from a group-level distribution, and it is possible to flexibly capture individual differences (*15*).

In addition to the ISS of resting-state FC, prior studies using diffusion-weighted imaging (DWI) have reported that white matter microstructure changes with aging (*16*). Structural connectivity (SC) reflects the interregional anatomical connections derived from DWI data, and FC is formed within the constraints of this anatomical architecture (*17, 18*). One previous study revealed that resting-state functional network integration appears to increase primarily when SC begins to decline (*19*).

This study aimed to identify the underlying factors of ISS decline during the Stroop task among older adults from the perspective of overactivation, using task-related ROIs derived from hierarchical Bayesian modeling, and to examine whether specific brain activation patterns are associated with task performance.

## 2. Results

### 2.1 Participant data

All older (N = 30) and younger participants (N = 22) conducted the Stroop task inside the MRI (Fig. 1A) and completed neuropsychological tests outside the MRI. One older adult with a Stroop task accuracy rate below 50% was excluded from the analysis. Participant characteristics and the behavioral performance of the Stroop task results are shown in Table 1. Neuropsychological tests revealed that the older group had significantly lower Mini-Mental State Examination (MMSE scores than the younger group (p < 0.05). The older group exhibited significantly longer reaction times than the younger group in both the congruent and incongruent conditions (p < 0.05). In contrast, no significant group differences were observed in the accuracy rates (congruent, p = 0.33; incongruent, p = 0.59). Furthermore, the mean interference index ΔRT, defined as the difference between reaction time in the incongruent and the congruent condition, was larger in older adults than that in younger adults but this difference did not reach statistical significance (p = 0.05).

**Figure 1.**
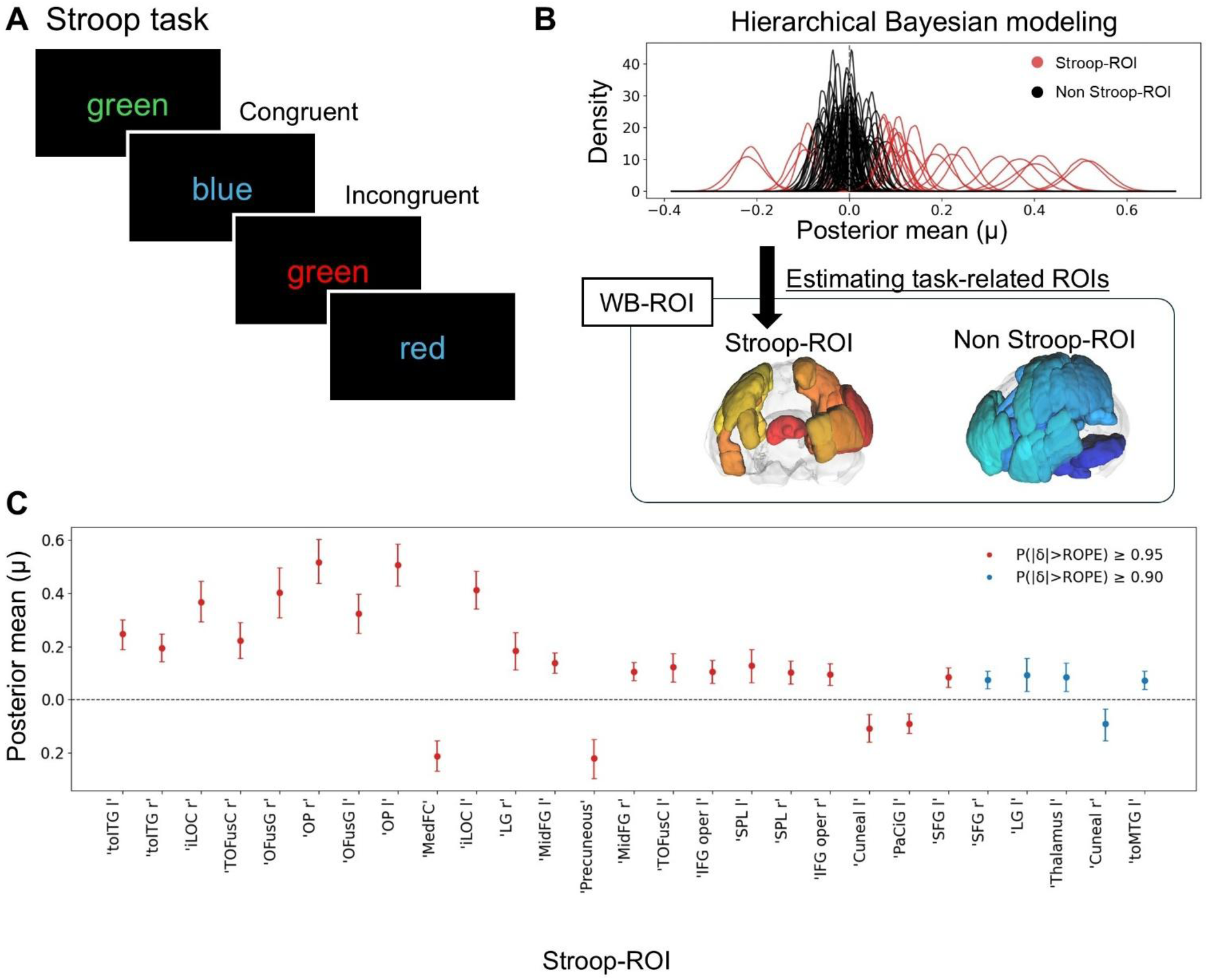
Identification of task-related brain regions using hierarchical Bayesian modeling. (A) Schematic of the Stroop task. Participants were presented with color words printed in congruent or incongruent ink colors and responded to the ink color. (B) Hierarchical Bayesian modeling was conducted to estimate task-related activation for each of 105 ROIs. The density plot shows the distribution of posterior means across all ROIs. ROIs with significant activation were designated as Stroop-ROIs (27 regions), and the remaining 78 ROIs were classified as non-Stroop-ROIs. (C) Posterior means of task-related activation for the 27 Stroop-ROIs. Error bars represent 95% highest density intervals. Red and blue points indicate ROIs meeting the credibility thresholds. The 27 Stroop-ROIs includ: inferior temporal gyrus, temporooccipital part left (toITG l), inferior temporal gyrus, temporooccipital part right (toITG r), lateral occipital cortex, inferior division right (iLOC r), temporal occipital fusiform cortex right (TOFusC r), occipital fusiform gyrus right (OFusG r), occipital pole right (OP r), occipital fusiform gyrus left (OFusG l), occipital pole left (OP l), frontal medial cortex (MedFC), lateral occipital cortex, inferior division left (iLOC l), lingual gyrus right (LG r), middle frontal gyrus left (MidFG l), precuneous cortex (Precuneous), middle frontal gyrus right (MidFG r), temporal occipital fusiform cortex left (TOFusC l), inferior frontal gyrus, pars opercularis left (IFG oper l), superior parietal lobule left (SPL l), superior parietal lobule right (SPL r), inferior frontal gyrus, pars opercularis right (IFG oper r), cuneal cortex left (Cuneal l), paracingulate gyrus left (PaCiG l), superior frontal gyrus left (SFG l), superior frontal gyrus right (SFG r), lingual gyrus left (LG l), thalamus left (Thalamus l), cuneal cortex right (Cuneal r), middle temporal gyrus, temporooccipital part left (toMTG l).

**Table 1.** Characteristics and group comparison of Stroop task performance.

|  | Young (n=22) | Old (n=29) | P value |
| --- | --- | --- | --- |
| Age (years) | 23.77 ± 5.23 | 68.48 ± 5.59 | - |
| Gender (Male/Female) | 15 / 7 | 14 / 15 | - |
| MMSE score | 29.54 ± 0.67 | 28.34 ± 1.70 | p < 0.05 |
| Stroop performance |  |  |  |
| Congruent RT (ms) | 0.66 ± 0.08 | 0.73 ± 0.07 | p < 0.01 |
| Incongruent RT (ms) | 0.69 ± 0.09 | 0.78 ± 0.08 | p < 0.01 |
| Δ RT (ms) | 0.03 ± 0.03 | 0.05 ± 0.03 | p = 0.05 |
| Congruent accuracy | 0.98 ± 0.02 | 0.98 ± 0.03 | p = 0.33 |
| Incongruent accuracy | 0.97 ± 0.03 | 0.94 ± 0.18 | p = 0.59 |

### 2.2 Identification of Stroop-related regions

Whole-brain activity was estimated using hierarchical Bayesian modeling to identify regions showing significant task-related activation (Fig. 1B). Significant activation ( μ_r_) was observed in 27 regions in both the younger and older groups (Fig. 1C). These 27 ROIs were defined as the Stroop-ROIs. The largest overactivation was observed in occipital-visual regions, including left lingual gyrus (LG) (δ = +0.243), left occipital fusiform gyrus (OFusG) (δ = +0.235), and right LG (δ = +0.204) (Supplementary Fig. 1).

### 2.3 Inter-subject similarity analysis

ISS analysis was conducted to evaluate the similarities in brain activity patterns. ISS analysis was performed using 105 whole-brain ROIs (WB-ROIs), including 78 non-Stroop-ROIs and 27 Stroop-ROIs (Fig. 2A). For each participant, the mean correlation coefficient with other participants within the same group was defined as the individual’s mean ISS (mISS). Remarkably, mISS in Stroop ROIs in older adults significantly decreased during task rather than younger adults (mISS = 0.61 ± 0.04 and 0.52 ± 0.05 for younger and older adults respectively, d = 1.88, p < 0.01) (Fig. 2B), but the mISS in the Stroop-ROIs showed no significant difference between younger (mISS = 0.62 ± 0.05) and older adults (mISS = 0.61 ± 0.04) at the resting state (d = 0.24, p = 0.44). In contrast, WB-ROI showed significant group difference in both resting state (young; mISS = 0.52 ± 0.04, old; mISS = 0.48 ± 0.04, d = 1.00, p < 0.01) and task condition (young; mISS = 0.50 ± 0.04, old; mISS = 0.44 ± 0.04, d = 1.53, p < 0.01). These results indicate that a task-specific functional change of the ISS decline in the Stroop-ROIs is independent of the general age-related alterations observed at resting state.

**Figure 2.**
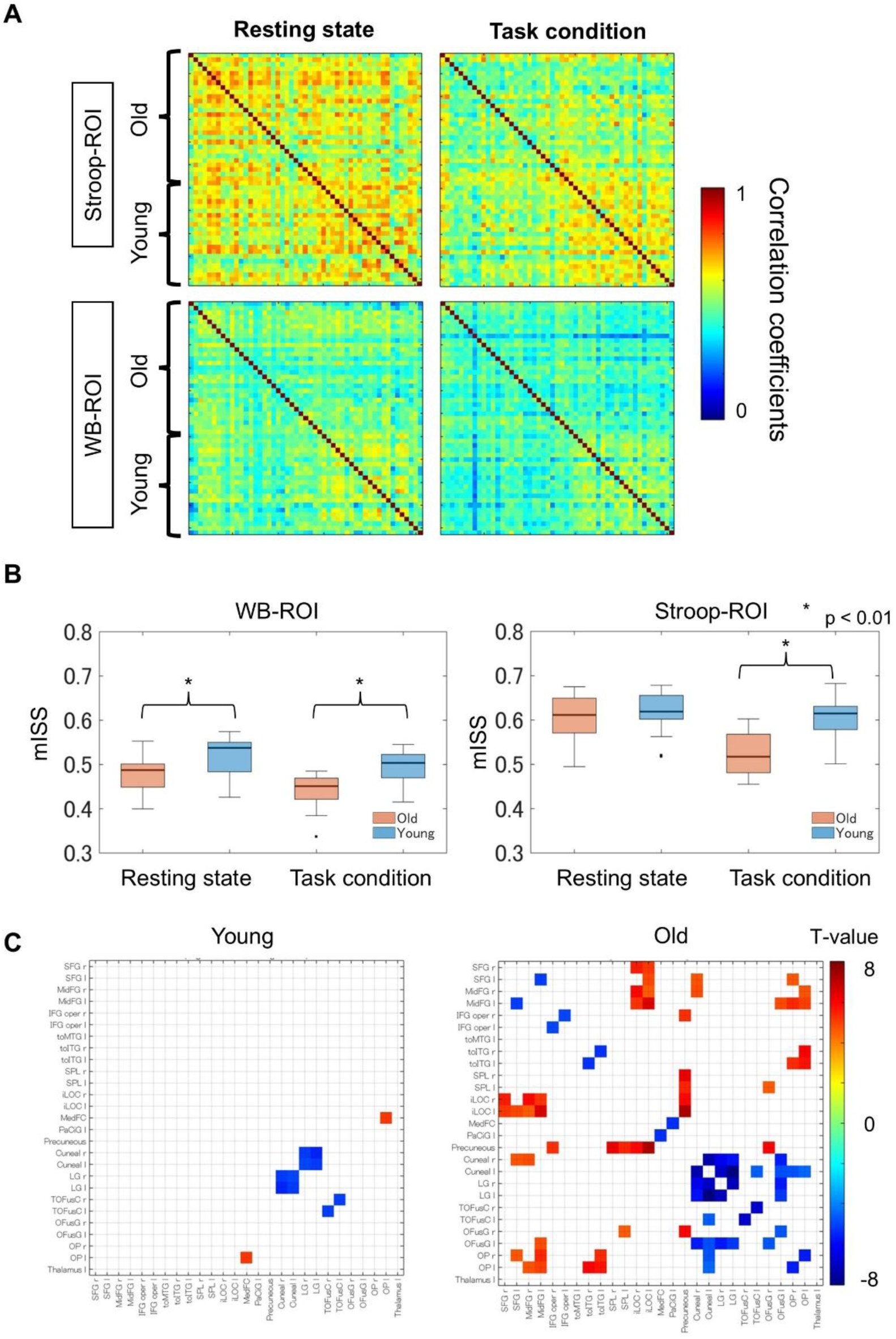
Inter-subject similarity (ISS) analysis. (A) ISS matrices showing pairwise correlation coefficients of functional connectivity (FC) feature vectors across all participants during resting state and task conditions. Matrices are shown separately for Stroop-ROIs and WB-ROIs. (B) Comparison of mean ISS (mISS) between younger and older groups for WB-ROIs and Stroop-ROIs during resting state and task conditions. (C) FC modulation from rest to task within the Stroop-ROIs for the younger and older groups. Color indicates t-values from one-sample t-tests on the FC difference. Only ROI pairs surviving Bonferroni correction are shown.

We also investigated the difference in FC with the Stroop-ROIs between the resting-state and task conditions (Fig. 2C). A few ROI pairs exhibited significantly altered FC in the younger group. In contrast, the older group showed extensive changes in FC. Interestingly, the connectivity strength decreased within the frontal and occipital lobes, whereas FC between the frontal and occipital lobe regions showed a significant increase. The differences in FC across the ROIs other than the Stroop-ROIs are presented in Supplementary Fig. 2. Several significant decreases in FC were observed in the younger group, and the older group exhibited a more widespread FC decrease. Compared with the Stroop-ROIs, fewer FCs showed a significant increase, indicating that the FC increase in older adults is specific to the Stroop-ROIs.

### 2.4 Relationship between group differences and individual variability

Pearson correlations between group difference parameter δ_r_ and ROI-specific standard deviation σ_a,r_ were computed. A significant positive correlation between δ_r_ and σ_a,r_ was observed across the Stroop-ROIs (r = 0.49, p < 0.01). However, scaling analyses revealed that this relationship was partially confounded by the mean-variance scaling effect. The correlation between task-related activation μ_r_ and σ_a,r_ was significant (r = 0.58, p < 0.01), indicating that ROIs with higher mean activation tended to have proportionally larger variance. When this scaling effect was controlled via partial correlation, the relationship decreased (r = 0.39, p < 0.05). Furthermore, when variability was expressed as the coefficient of variation, the correlation with δ was eliminated (r = −0.001, p = 0.995).

### 2.5 Leave-one-out ROI analysis

We then conducted a leave-one-out ISS analysis to assess the causal contribution of each ROI to the ISS group difference. The Δd was defined as the difference between the effect size of the group difference across all Stroop-ROIs and that obtained after excluding a specific ROI, and was correlated with σ_a,r_ and δ_r_. A significant negative correlation between σ_a,r_ and Δd (r = −0.59, p < 0.01) was observed (Supplementary Fig. 3). Notably, δ_r_ showed no significant correlation with Δd (r = −0.17, p = 0.40). The regions increasing the ISS group difference most strongly were the frontal areas (Fig. 3A), including the left middle frontal gyrus (MidFG) (Δd = +0.30), right MidFG (Δd = +0.23), left inferior frontal gyrus, pars opercularis (IFGoper) (Δd = +0.20), and right IFGoper (Δd = +0.20). In contrast, the regions suppressing the ISS group difference was the occipital-visual areas, including the left inferior division of lateral occipital cortex (iLOC) (Δd = −0.29) and right OFusG (Δd = −0.25). Group-specific decomposition revealed that these two sets of regions were derived from contributions from different groups (Fig. 3B). The frontal regions contributed to the ISS group difference by maintaining high FC consistency in the younger group, whereas the occipital-visual regions buffered it by maintaining FC consistency in the older group (Fig. 3C).

**Figure 3.**
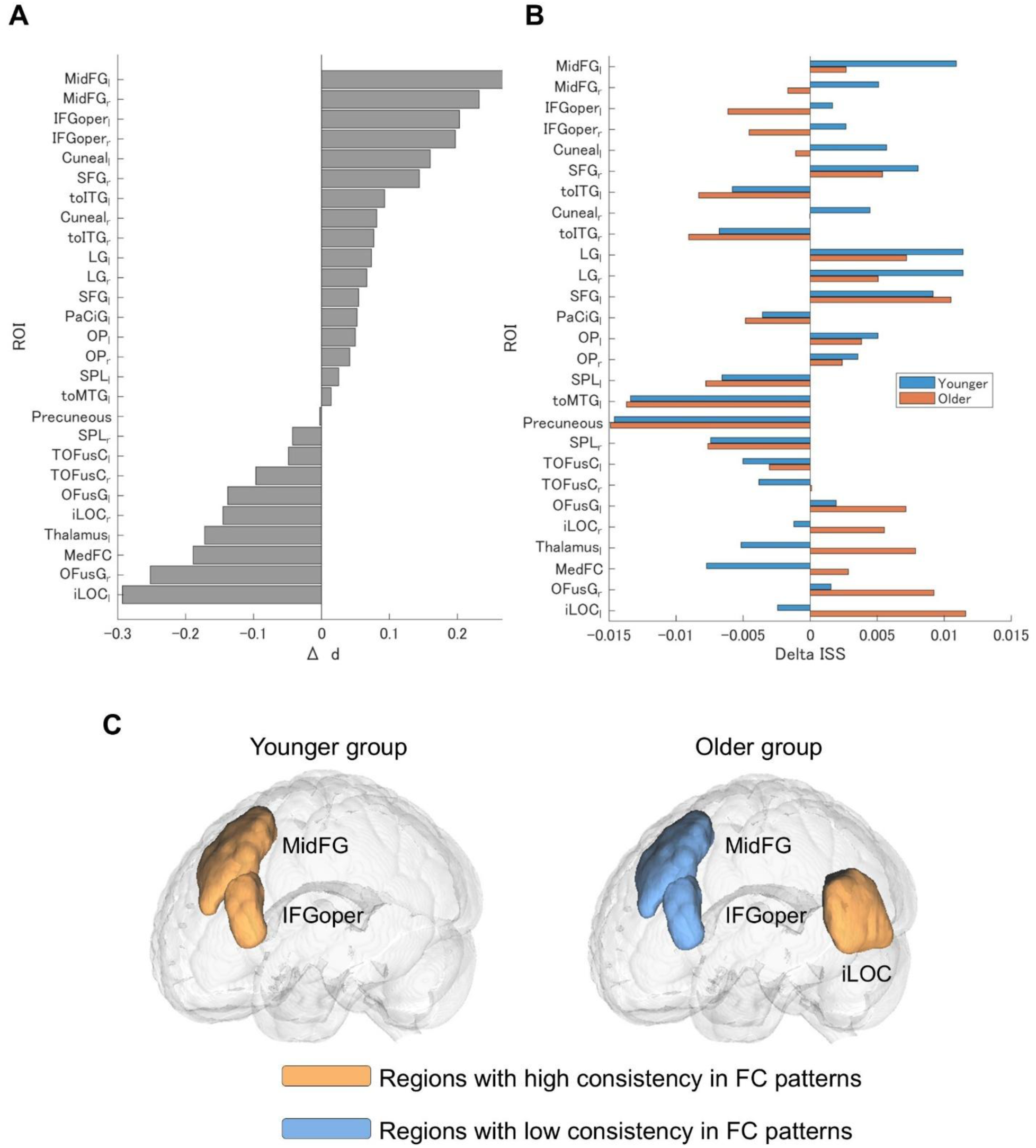
Leave-one-out ROI analysis reveals distinct regional contributions to ISS group differences. (A) Change in effect size of the ISS group difference (Δd) upon exclusion of each Stroop-ROI. Positive Δd indicates that the ROI increases the ISS group difference, whereas negative Δd indicates the ROI suppresses it. (B) Group-specific decomposition of the leave-one-out effect (ΔISS) for each ROI, shown separately for younger (blue) and older (orange) groups. Positive ΔISS indicates that the excluded ROI contributed to within-group FC consistency. (C) Brain surface rendering illustrating the spatial distribution of regions with high (orange) and low (blue) FC pattern consistency in younger and older groups.

### 2.6 Principal component analysis of activation patterns

The hierarchical Bayesian model provided the a_i,r_ matrix, where each element represents the subject-specific deviations in activation. Principal component analysis (PCA) applied to the a_i,r_ matrix of the older group yielded principal component 1 (PC1) accounting for 72.9% of the variance, PC2 accounting for 6.8%, and PC3 accounting for 5.4%. Only PC3 showed a significant positive correlation with ΔRT (r = 0.48, Bonferroni-corrected P < 0.05) (Fig. 4A). In PC3, ROIs with the large positive loadings were the precuneus (+0.43), frontal medial cortex (MedFC; +0.37), left thalamus (+0.30), and paracingulate gyrus (PaCiG; +0.21), whereas those with the largest negative loadings were the iLOC (−0.37), occipital pole (OP; −0.26), and superior parietal lobule (SPL; −0.22) (Fig. 4B). Together, older adults whose activation was biased towards the medial default mode network (DMN) regions tended to exhibit larger ΔRT values, indicating poorer performance. In contrast, those whose activation was biased toward visual attention regions tended to exhibit smaller ΔRT values, indicating better performance. PC1, which represents variation in overall activation level (Supplementary Fig. 4), showed no significant correlation with ΔRT (r = −0.07, p = 0.73), indicating that global variations in activation levels were unrelated to Stroop performance.

**Figure 4.**
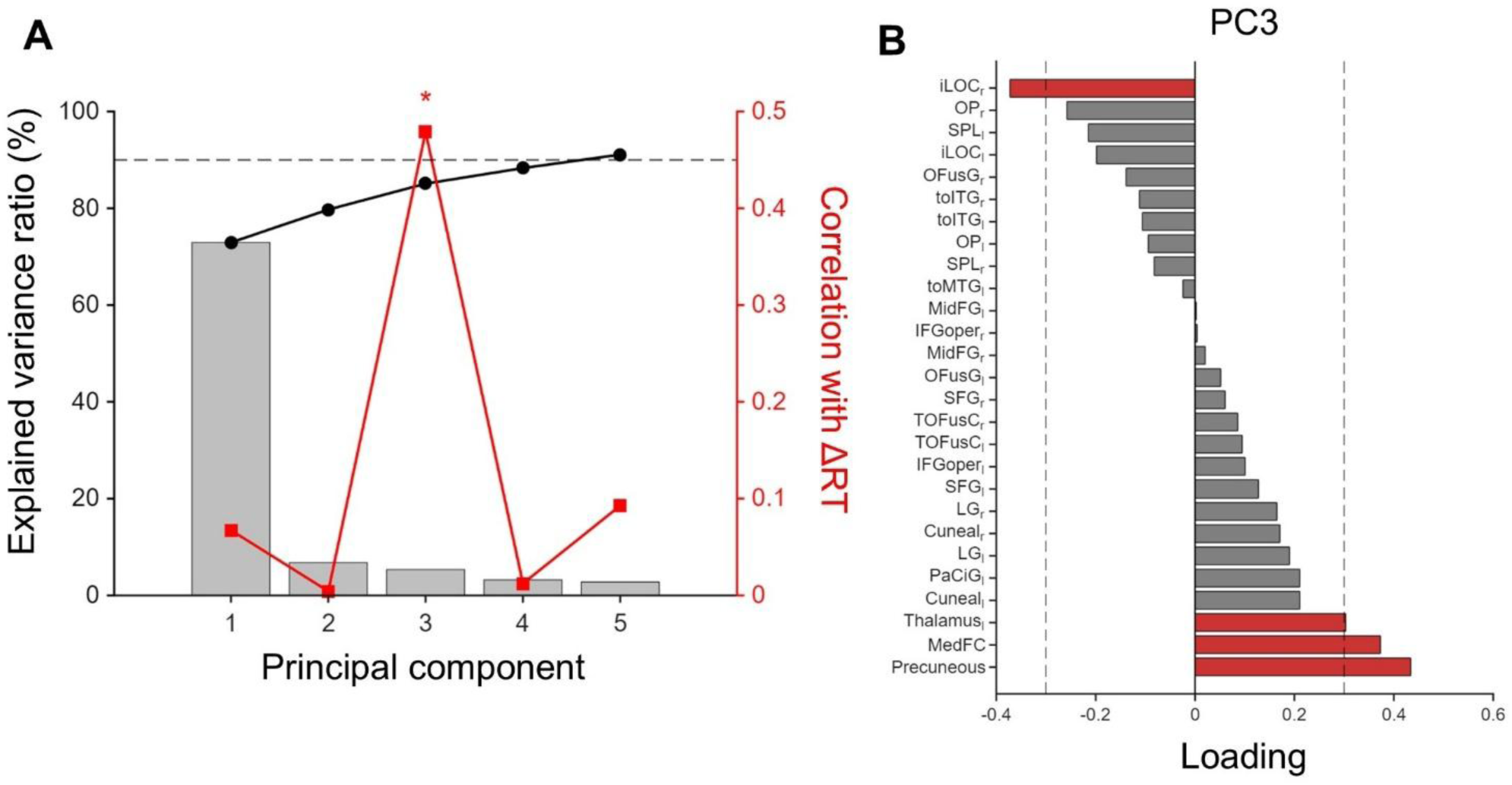
Principal component analysis identifies an activation pattern linked to Stroop performance in older adults. (A) Explained variance ratio (gray bars) and correlation with ΔRT (red points) for each principal component derived from the subject-specific activation deviation matrix of the older group. The dotted line represents the 90% cumulative variance threshold. (B) Factor loadings of PC3 across the Stroop-ROIs. Factor loadings with absolute values of 0.3 or greater are highlighted in red.

### 2.7 Structural connectivity

SC analysis was conducted to verify whether changes in FC were attributable to the anatomical degeneration of nerve fibers. Edgewise group comparisons across all 5,460 connections revealed 213 edges with significantly lower SC in the older group (p < 0.05, FDR-corrected) (Fig. 5A). The Stroop-ROI pairs were assessed using enrichment analysis to evaluate the concentrated age-related SC decreases. Three connectivity categories were defined: Stroop-ROI pairs, pairs of the Stroop-ROIs and non-Stroop-ROIs, and pairs of non-Stroop-ROIs (Fig. 5A). An enrichment ratio greater than 1 indicates that significant SC decreases are concentrated in that category. Only four connections among the Stroop-ROI pairs showed significant SC reduction (p < 0.05, FDR-corrected), yielding an enrichment ratio of 0.34. This was substantially lower than the rate observed among non-Stroop-ROI pairs (enrichment ratio = 1.19), indicating that age-related white matter degeneration tends to occur within non-Stroop-ROIs. Furthermore, no significant correlations were found between SC strength in the hub regions and ΔRT (Fig. 5B).

**Figure 5.**
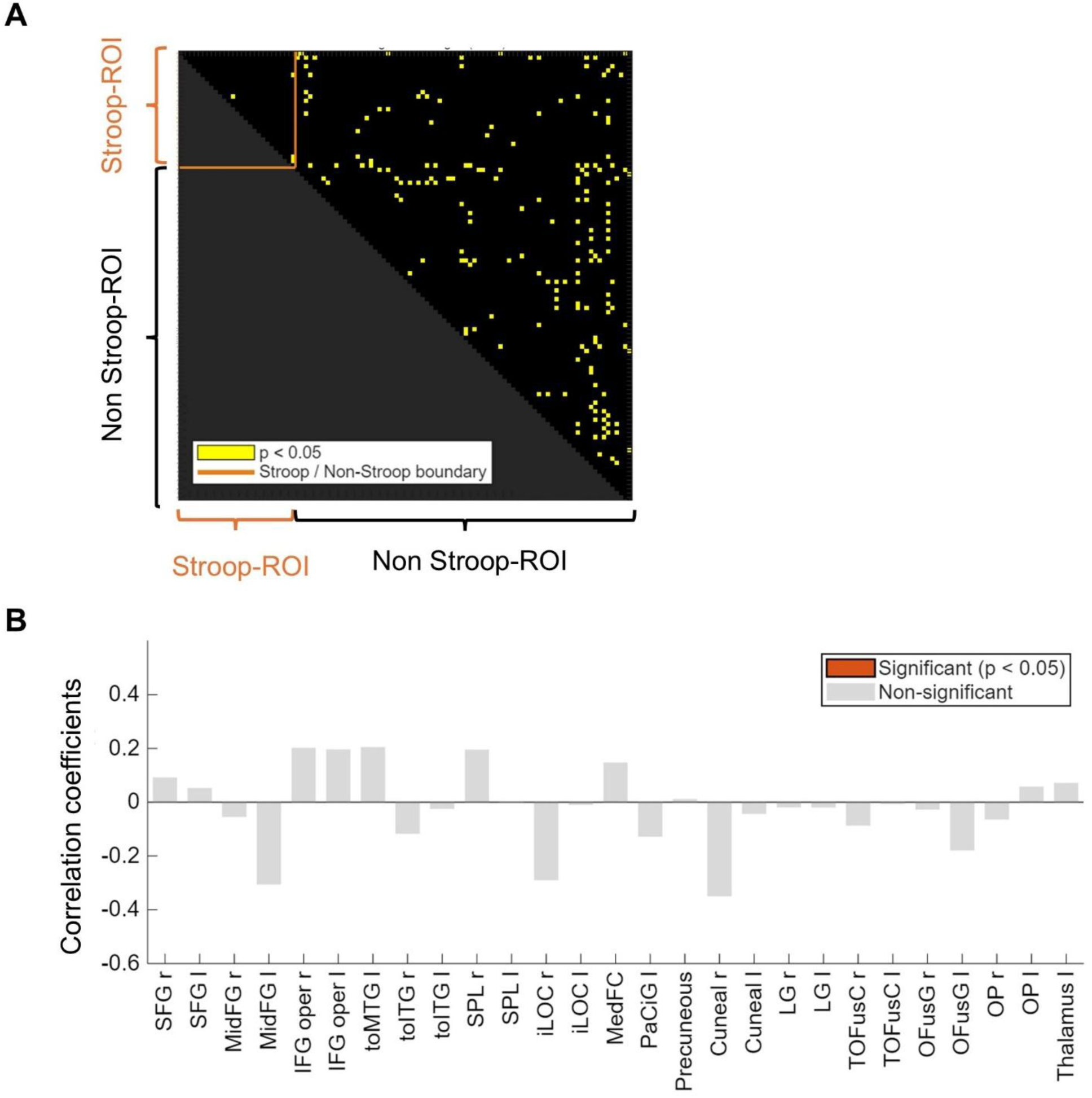
Structural connectivity (SC) analysis. (A) SC matrix showing edge-wise group comparisons between older and younger group. Yellow points indicate connections with significantly lower SC in the older group (p < 0.05, FDR-corrected). The orange line represents the boundary between Stroop-ROIs and non-Stroop-ROIs. (B) Correlations between SC strength and ÄRT for each Stroop-ROI region. No significant associations were observed.

## 3. Discussion

This study clearly demonstrates not only factors underlying ISS reduction during performing the Stroop task in older adults from the perspective of overactivation, but also the relationship between brain activity patterns and task performance. We first, using hierarchical Bayesian estimation, identified the Stroop-ROIs and, importantly, these Stroop-ROIs includes the several brain regions related to the cognitive inhibition (*20*). The WB-ROIs showed significant ISS group differences both task condition and resting state. In contrast, the Stroop-ROIs showed significant ISS group differences during the task condition only. These observations indicate that the ISS decline in task-related regions reflects functional change related to the task, rather than a general age-related deterioration already present in the resting state.

The combination of hierarchical Bayesian estimation and leave-one-out ISS analysis showed that frontal regions maintain high FC patterns consistency in younger group but not in the older group. In contrast, occipital regions maintain FC consistency in the older group. Furthermore, by using the individual deviation matrix a_i,r_ derived from this model as the input for PCA, specific brain activity patterns associated with Stroop task performance was identified.

### 3.1 Identification of Stroop-related regions

In this study, we identified 27 Stroop-ROIs using a hierarchical Bayesian model (Fig. 1C). These regions included multiple brain areas involved in the Stroop task. The IFGoper is involved in suppressing prepotent responses (*21*), the MidFG is involved in attentional control and conflict processing (*22*). The PaCiG is involved in monitoring response conflict (*23*), and the superior frontal gyrus has been reported to be associated with higher-order cognitive control, including inhibitory control (*24*). Furthermore, we observed the occipital visual areas, including OFusG, iLOC, LG, occipitotemporal gyrus, and temporo-occipital fusiform cortex. Although these regions are not directly involved in cognitive inhibition, color-word conflict processing is essential in the Stroop task, and these areas were likely extracted as a visual-processing foundation supporting task performance. In addition, regions such as the precuneus, MedFC, and thalamus were also included, indicating that the Stroop-ROIs captured not only core regions of cognitive inhibition, but also widespread neural activity associated with task.

### 3.2 Task specificity of ISS reduction

In the Stroop-ROIs, no significant difference in ISS was observed during the resting state, while significant group differences were found during task performance. In contrast, the WB-ROIs showed group differences in both the resting state and task condition (Fig. 2B). These results indicate that ISS reduction in the Stroop-ROIs reflects the functional change that occurs during task performance, rather than a general age-related change at resting state. Previous studies have reported whole-brain level ISS reduction in older adults, which becomes more pronounced during task performance (*25, 26*). Furthermore, a detailed examination of the FC change patterns revealed widespread FC decreases within the frontal and occipital regions from rest to task but fronto-occipital FC increased in the older group (Fig. 2C). This widespread changes in FC were not observed in the younger group. These results indicate the intermodule integration increases during task performance in older adults, supported by previous study (*27*). The enhancement of frontal-occipital connectivity possibly reflects a part of this age-related network change.

### 3.3 Determinants of ISS reduction

If overactivation reflects the noise, regions with greater overactivation should exhibit greater inter-individual variability and this variability should lead to ISS reduction. Given the positive correlation between the magnitude of overactivation ( δ_r_) and inter-individual variability across ROIs (σ_a,r_) (r = 0.49) was was observed, it was considered that these factors might reflect the noise. However, this association could be partially explained by the statistical properties of mean-variance scaling, whereby variance increases proportionally with the mean. When normalized by the coefficient of variation, the correlation with δ_r_ disappeared (r = −0.001). These findings suggest that the association between overactivation and inter-individual variability is more likely explained by mean-variance scaling, rather than indicating that overactivation reflects noise.

Furthermore, the results of the leave-one-out ISS analysis showed that σ_a,r_ was significantly correlated with the contribution to group differences in ISS (Δd) (r = −0.59), whereas δ_r_ did not correlate with Δd (r = −0.17). This indicates that smaller variability in regional activity is associated with a larger contribution to the difference in ISS between younger and older groups. In addition, the magnitude of overactivation ( δ_r_) was not significantly correlated with Δd (r = −0.17). The regions driving ISS reduction primarily included the frontal regions, such as the MidFG and IFGoper (Fig. 3A); conversely, the occipital regions, such as the iLOC and OFusG, which show large inter-individual variability, suppressed ISS group differences. This result represents the opposite pattern from the noise model that “regions with greater variability drive ISS reduction”. We therefore conclude that the ISS reduction from the perspective of overactivation is not due to “noise” but to “compensation” for functional decline to maintain cognitive performance.

The leave-one-out analysis was also performed within each group, and ΔISS, defined as the difference between the mISS across all 27 ROIs and the mISS with a specific ROI excluded, was calculated for both the younger and older groups. In the MidFG and IFGoper, ΔISS was larger in the younger group, and smaller in the older group (Fig. 3B). This indicates that frontal regions support FC consistency in younger group; however, this supportive function was decreased in older group. This finding reflects the loss of consistency in frontal regions with aging, which is consistent with previous findings on age-related changes in frontal function (*28, 29*). Conversely, excluding the iLOC reduced ISS in the older group, whereas the effect in the younger group was small (Fig. 3B). This indicates that the iLOC helps to maintain FC consistency in older group, thereby reducing ISS group differences. In other words, overactivation and large interindividual variability at the activation level do not necessarily signify a breakdown of the network structure. Instead, ISS reduction is determined by the functional reduction of the frontal network, which maintains high consistency in younger group, and the emergence of the occipital network, which contributes to maintaining consistency in older group.

### 3.4 Principal component analysis of activation patterns

PCA reveals that Stroop task performance (ΔRT) in the older adult group can be predicted not by the variation in overall activation level (PC1), but by a specific spatial pattern (PC3) (Fig. 4A). Previous studies have shown that the first principal component in fMRI data is almost entirely equivalent to the global signal (*30*). PC3 reflected the contrast between the precuneus, MedFC, and left thalamus (positive loadings), as well as the iLOC, OP, and SPL (negative loadings), indicating that older adults with an increased DMN activity were negatively correlated with performance, whereas posterior visual and dorsal attention activity were positively associated with performance. Additionally, iLOC was a region that showed significantly higher activity in older group than in younger group (Supplementary Fig. 1). These results indicate that overactivation in older adults is not a uniform phenomenon, and that behavioral consequences depend on the spatial distribution of activation. Furthermore, the loading pattern of PC3 can be interpreted as a failure of DMN deactivation during task performance. Previous studies have similarly reported that DMN suppression during task performance is insufficient in older adults (*31, 32*); further, it has been suggested that this deactivation failure causes resource competition with task processing (*33*). These results are consistent with the present study. Although PC3 explained only 5.4% of the variance, it was significantly associated with behavior, indicating that age-related brain activity changes influence behavior through specific spatial contrasts, rather than large variance components.

### 3.5 Structural connectivity

Although significant SC reductions were observed across 213 edges at the whole-brain level (Fig. 5A), they were not concentrated in connections within the Stroop-ROIs (enrichment ratio, 0.34). Furthermore, no significant correlation was identified between the SC strength of each region in the Stroop-ROI and ΔRT (Fig. 5B). Prior studies have reported that white matter structural changes with aging are associated with declines in executive function (*34*); however, the Stroop-ROIs extracted in this study showed relatively few age-related effects. These results suggest that FC changes and ISS reduction in the Stroop-ROIs reflect functional, rather than anatomical changes.

## 4. Conclusion

We clearly showed that brain regions related to performing Stroop task behave differently between the younger and older adults in task condition but not in resting state. The hierarchical Bayesian model is a better approach for Stroop-ROI estimation, as it allows for individual variation while coherently pooling information across individuals, compared with the general linear model. Stroop task performance in older adults was associated not with overall activation levels, but rather with a specific spatial pattern reflecting the balance between DMN and visual attention regions. These findings suggest that such mechanisms play an important compensatory role in age-related functional decline.

## 5. Methods

### 5.1 Participants

This study was conducted with 30 healthy older adults (mean age 68.5 ± 5.5 years, 15 males, 15 females) and 22 young adults (mean age 23.8 ± 5.2 years, 15 males, 7 females). Participants were recruited via online bulletin boards, and data were collected from October 2024 to October 2025. Age groups were defined as young adults (18–39 years old) and older adults (60–79 years old). The inclusion criteria were individuals with the ability to provide written informed consent for voluntary participation in the study, aged between 18 and 80 years at the time of consent, and native Japanese speakers. Exclusion criteria were as follows: pregnant women, individuals with higher brain dysfunction or psychiatric disorders, individuals with visual impairments, individuals requiring assistance with walking, individuals with a history of cerebrovascular or cardiovascular disease, individuals with metal, medical devices, or implants in the body, individuals with tattoos, individuals with claustrophobia, individuals weighing 100 kg or more. All participants underwent the MMSE and MRI. Additionally, the Trail Making Test (TMT) A and B, Timed Up and Go test (TUG), and dual-task TUG were used for other experiments. This study was approved by the Ethics Committee of the National Institute of Advanced Industrial Science and Technology (AIST).

### 5.2 MRI acquisition

All MRI experiments were conducted using a 3 Tesla MRI scanner (Ingenia, Philips) equipped with a 32-channel head coil. Imaging sequences included fMRI during Stroop task execution, resting state fMRI, structural images, and diffusion-weighted imaging (DWI). For Stroop task fMRI, each session consisted of an initial 10-second rest period followed by alternating 30 second task blocks and 30 second rest blocks presented five times, with a total of two sessions performed. During task blocks, one of three Japanese words (“red”, “green”, or “blue”) was presented every 2 seconds, and participants were instructed to respond to the ink color rather than the word meaning. Responses were made using button presses with the right hand, selecting the correct color from three buttons corresponding to each color. The difference between the mean reaction time for incongruent trials and that for congruent trials (ΔRT) was calculated as an index of each participant’s performance on the Stroop task. Each task block included congruent conditions where word meaning and color matched and incongruent conditions, with stimulus presentation order randomized. Participants practiced the Stroop task outside the MRI room beforehand to ensure adequate understanding before scanning. Task presentations used PsychoPy (*35*), and accuracy and response times were recorded. Stroop task fMRI parameters were as follows: T2*-weighted gradient-echo echo planner imaging (EPI), time repetition (TR)/ echo time (TE) = 1,500/30 ms, voxel size = 2.5 × 2.5 mm², slice thickness = 2.5 mm, matrix size = 76 × 76, 44 slices, flip angle = 80°, multiband (MB) factor = 2, 210 scans. Resting state fMRI parameters were as follows: T2*-weighted gradient-echo EPI, TR/TE = 1,500/30 ms, voxel size = 2.5 × 2.5 mm², slice thickness = 2.5 mm, matrix size = 76 × 76, 44 slices, flip angle = 80°, MB factor = 2, 420 scans. T1 parameters were as follows: magnetization prepared rapid acquisition gradient echo sequence for registration in preprocessing TR/TE = 11.1/5.1 ms, voxel size = 0.7 × 0.7 × 0.7 mm³, matrix size = 365 × 342 × 257. DWI parameters were as follows: diffusion weighted spin echo EPI, TR = 7,950 ms, TE = 95 ms, voxel size = 2.5 × 2.5 × 2.5 mm³, matrix size = 72 × 72 × 52, b-value = 0, 1000, 3000 s/mm², 64 directions, MB factor = 4 and images with reversed phase-encoding direction at b = 0 were acquired for distortion correction.

### 5.3 MRI data preprocessing

One older adult with a Stroop task accuracy rate below 50% was excluded from the analysis, leaving 29 older and 22 younger adults. MRI data preprocessing was conducted using SPM12 (Wellcome Trust Center for Neuroimaging, UK). The preprocessing steps included slice-timing correction, realignment for motion correction, and normalization to the standard MNI space. Subsequently, the functional images were smoothed (FWHM kernel: 7.5 × 7.5 × 7.5 mm³). For the fMRI data collected during the Stroop task, a general linear model (GLM) analysis was performed for each participant to detect task-related brain activity. Six head motion parameters were included as covariates in the model. In this study, regressors were created for each task block, generating β maps of task-related brain activity for 10 blocks per participant. The β values indicate the magnitude of contribution of each task block regressor to the blood oxygenation level dependent (BOLD) signal, and serve as an index of task-related BOLD response amplitude. Subsequently, the mean β values for each region of interest (ROI) were calculated based on the Harvard-Oxford Atlas implemented in the CONN toolbox (*36*). The ROIs included 91 cortical and 14 subcortical regions in both hemispheres. Finally, 105 ROI mean β values were calculated from 10 β maps for each subject.

### 5.4 Hierarchical Bayesian model

The β values estimated for each subject, each block, and each ROI were used as observed data. For group comparison, age group variables were centered, with the young group labeled −0.5 and the older group labeled 0.5, such that the intercept term represented the average activity level of both groups. Analysis used the Python library PyMC version 5. Input data consisted of a matrix containing subject information, group variables, task block information, ROI information, and β values. Each data point was defined as the observed beta value y_i,r_ for subject i and ROI r. Group information for subject i was represented as g_i_. For each ROI, we estimated the main effect parameter μ_r_ representing the task effect and the group difference effect δ_r_ representing the difference between young and older groups. Additionally, individual differences due to subject-ROI interactions were modeled as a_i,r_. These relationships were expressed in the following linear model:

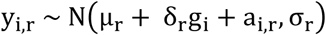

Laplace distributions were assigned as prior distributions for μ_r_ and δ_r_. Laplace distributions have the property of sparsification, shrinking estimates for ROIs with small effects toward zero. This automatically extracted ROIs with high activity. For the scale parameters b_μ_ and b_δ_ of the Laplace distributions, half-normal distributions were assigned as hyperpriors, allowing the strength of sparsification to be automatically estimated from the data.

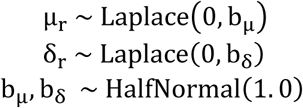

Regarding the random effect of subject by ROI a_i,r_, we assumed a normal distribution with different variances for each ROI. Specifically, this was expressed by multiplying a latent variable z_i,r_ following a standard normal distribution by a ROI-specific scale parameter σ_a,r_.

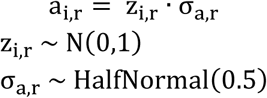

The standard deviation of observation error σ_r_ was also estimated independently for each ROI, with the following half-normal distribution assigned as a prior distribution.

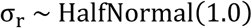

All parameter estimation in this study was performed using Markov Chain Monte Carlo (MCMC) method. Sampling conditions were set as follows: 4 chains, 3,000 tuning steps, 5,000 effective samples per chain, target acceptance of 0.95, and max tree depth of 14. The random seed was fixed in 2025 to ensure reproducibility. Convergence was assessed using the Gelman–Rubin statistic (R^). As an index of Bayesian credibility, probabilities P(|μ| > ROPE) and P(|δ| > ROPE) were calculated using region of practical equivalence (ROPE) (±0.05). The criterion for selecting significant activity regions was P(|μ| > ROPE) > 0.9.

### 5.5 Inter-subject similarity analysis

All fMRI data were processed using the CONN toolbox version conn22a. Six parameters of head motion and signals from the white matter and cerebrospinal fluid were regressed out, and detrending was applied to the residuals of the fMRI signal. For the resting-state fMRI data, low-frequency fluctuation components were extracted using a bandpass filter (0.008–0.09 Hz). Subsequently, ROI-to-ROI analysis was conducted using WB-ROIs and the Stroop-ROIs for both conditions during the Stroop task and resting state, creating FC matrices for each subject. The upper triangular components of each FC matrix were vectorized to generate the feature vectors for each subject. Correlation coefficients between the feature vectors across all subjects were calculated, and ISS analysis was employed. The mean correlation coefficient with other subjects within the same group was defined as the mISS. This procedure was applied separately during the Stroop task and in the resting state. The Mann–Whitney U test was used to compare the mISS between the younger and older groups. Cohen’s d was calculated as an index of the effect size. Analyses were conducted separately for the WB-ROIs and the Stroop-ROIs.

Within-group analyses were conducted to identify ROI pairs showing significant FC modulation from rest to task to characterize the FC changes underlying ISS group differences. For each participant, the difference between task condition and resting-state FC was computed for all unique ROI pairs. Within each age group, one-sample t-tests were conducted separately to identify ROI pairs with significant FC changes from rest to task. The Bonferroni correction was applied to control for multiple comparisons across all 351 ROI pairs.

### 5.6 Relationship between group differences and individual variability

The group difference parameter δ_r_ and the ROI-specific standard deviation σ_a,r_ for each ROI were derived from the hierarchical Bayesian model. Pearson correlations were computed between the δ_r_ and σ_a,r_ values. Because a larger mean activation can produce proportionally larger variance, we assessed the correlation between δ_r_ and the coefficient of variation, which normalizes the variability by mean activation level, and the partial correlation between δ_r_ and σ_a,r_ controlling for the overall mean activation μ_r_ to confirm the presence of a scaling effect.

### 5.7 Leave-one-out ROI sensitivity analysis

A leave-one-out ISS analysis, in which each ROI was excluded, was subsequently conducted. The change in the effect size of the group difference of ISS upon exclusion of each ROI (Δd) was computed and its Pearson correlation with σ_a,r_ and δ_r_ were calculated. Positive and negative Δd values indicate that the ROI increases and decreases the group difference of ISS, respectively. We also examined the group difference in variance and its correlation with Δd to assess whether group differences in neural variability contributed to the overall ISS group difference. Furthermore, we decomposed the leave-one-out effect for each ROI into group-specific contributions to clarify whether the ISS group difference was driven by changes in older, younger, or both groups. Specifically, we computed the change in the mean ISS for each group separately by subtracting the baseline mean ISS from the mean ISS value obtained after excluding each ROI. This was conducted independently for the older (ΔISS_OA) and younger (ΔISS_YA) groups. A positive ΔISS indicates that the excluded ROI contributed to within-group ISS consistency for that group, whereas a negative ΔISS indicates that the ROI reduced within-group consistency. By comparing the relative magnitudes of ΔISS_OA and ΔISS _YA, we determined whether each ROI’s contribution to the overall ISS group difference originated primarily from the older, younger, or both groups.

### 5.8 Principal component analysis of activation patterns

Using the activation deviations a_i,r_ estimated from the hierarchical Bayesian model, PCA was applied to the a_i,r_ matrix of the older adult group (29 participants × 27 ROIs). Pearson correlations were computed between the scores of each principal component (PC) accounting for up to 90% of cumulative variance and ΔRT. The Bonferroni correction was applied to control for multiple comparisons.

### 5.9 Structural connectivity analysis

One older adult with missing DWI data was excluded, resulting in a sample of 28 older and 22 younger adults. For analysis of SC, DWI data were preprocessed using MRtrix3 version 3.0.4 (*37*) and FSL (*38*). Preprocessing steps included denoising, distortion correction, eddy current correction, motion correction, B1 field inhomogeneity correction, and removal of non-brain tissue. ANTs version 2.5.0 (*39*) was used for registration to the template image. ROIs used were the same as 105 whole-brain regions as in FC analysis. After estimating fiber orientation distributions using MRtrix3, whole-brain probabilistic tractography was performed using the tckgen function (100 million streamlines). The resulting streamlines were optimized using the tcksift2 function, and a 105 × 105 SC matrix was generated for each participant, with each element representing the streamline count connecting a given ROI pair. Each element was normalized by the total number of streamlines in the whole brain. Edge-wise two-sample t-tests were performed on all connections to identify connections with significant age-related SC reduction. Multiple comparison correction was applied using false discovery rate (FDR) procedure. Enrichment analysis was performed by comparing the proportion of FDR significant edges. In addition, correlations between SC strength and ΔRT were examined for the hub regions identified by the FC analysis.

## Acknowledgments

We thank Ms. Mako Otobe for assistance of MRI experiment.

---

**Supplementary Figure 1.**
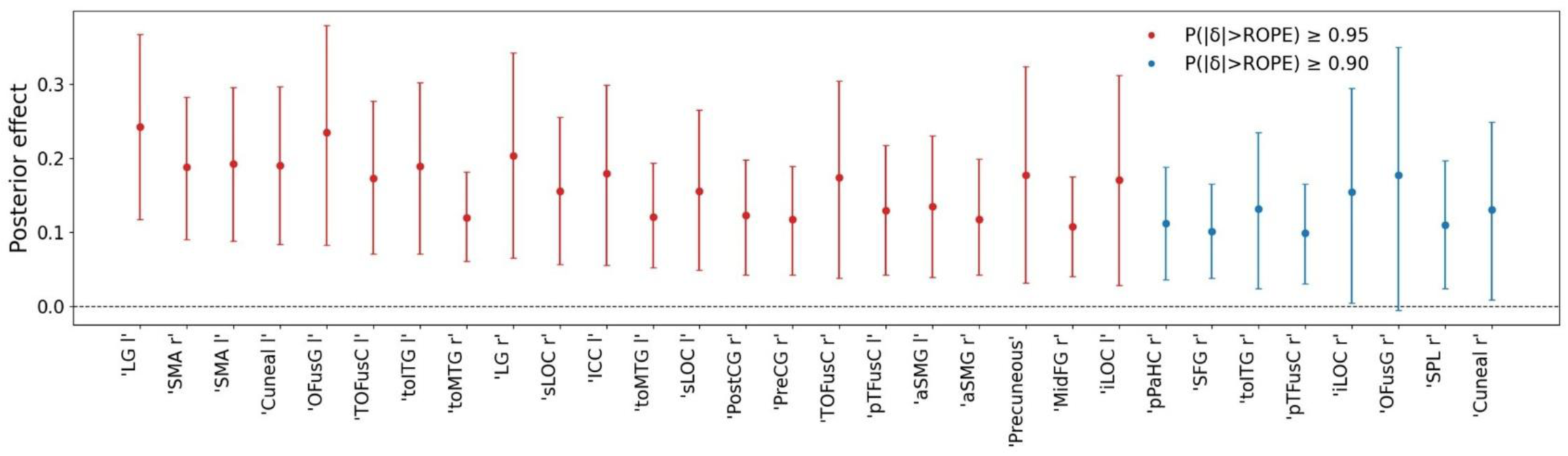
Estimated group difference parameters from the hierarchical Bayesian model. Posterior means of the group difference parameter estimated from the hierarchical Bayesian model. Positive values indicate brain regions where activity is higher in the older group compared to the younger group. Error bars represent 95% highest density intervals. Red and blue points indicate ROIs meeting the credibility thresholds.

**Supplementary Figure 2.**
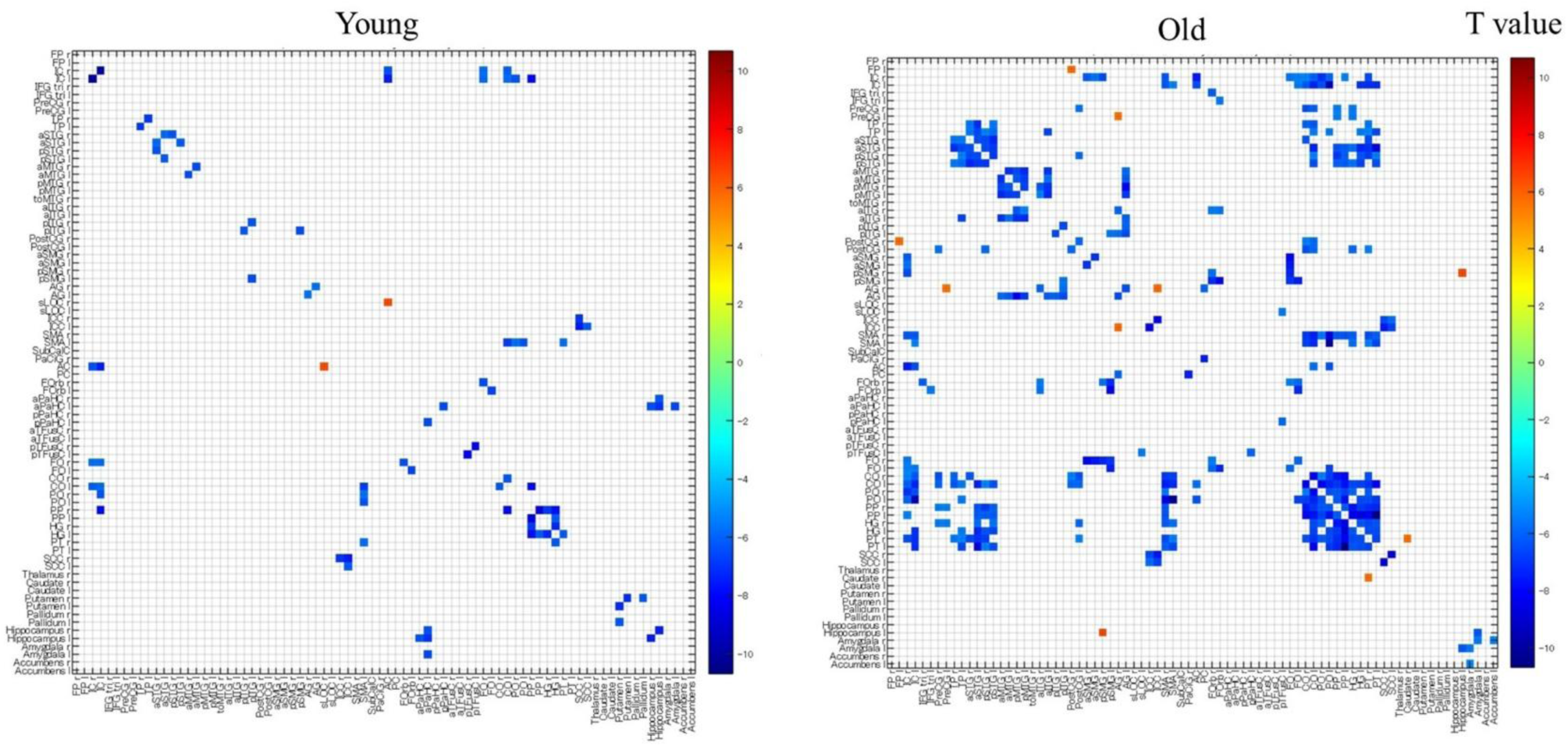
FC modulation from rest to task in non-Stroop-ROIs. FC differences between task condition and resting state for all ROI pairs involving non-Stroop-ROIs, shown separately for the younger and older groups. Color indicates t-values from one-sample t-tests. Only ROI pairs surviving Bonferroni correction are displayed.

**Supplementary Figure 3.**
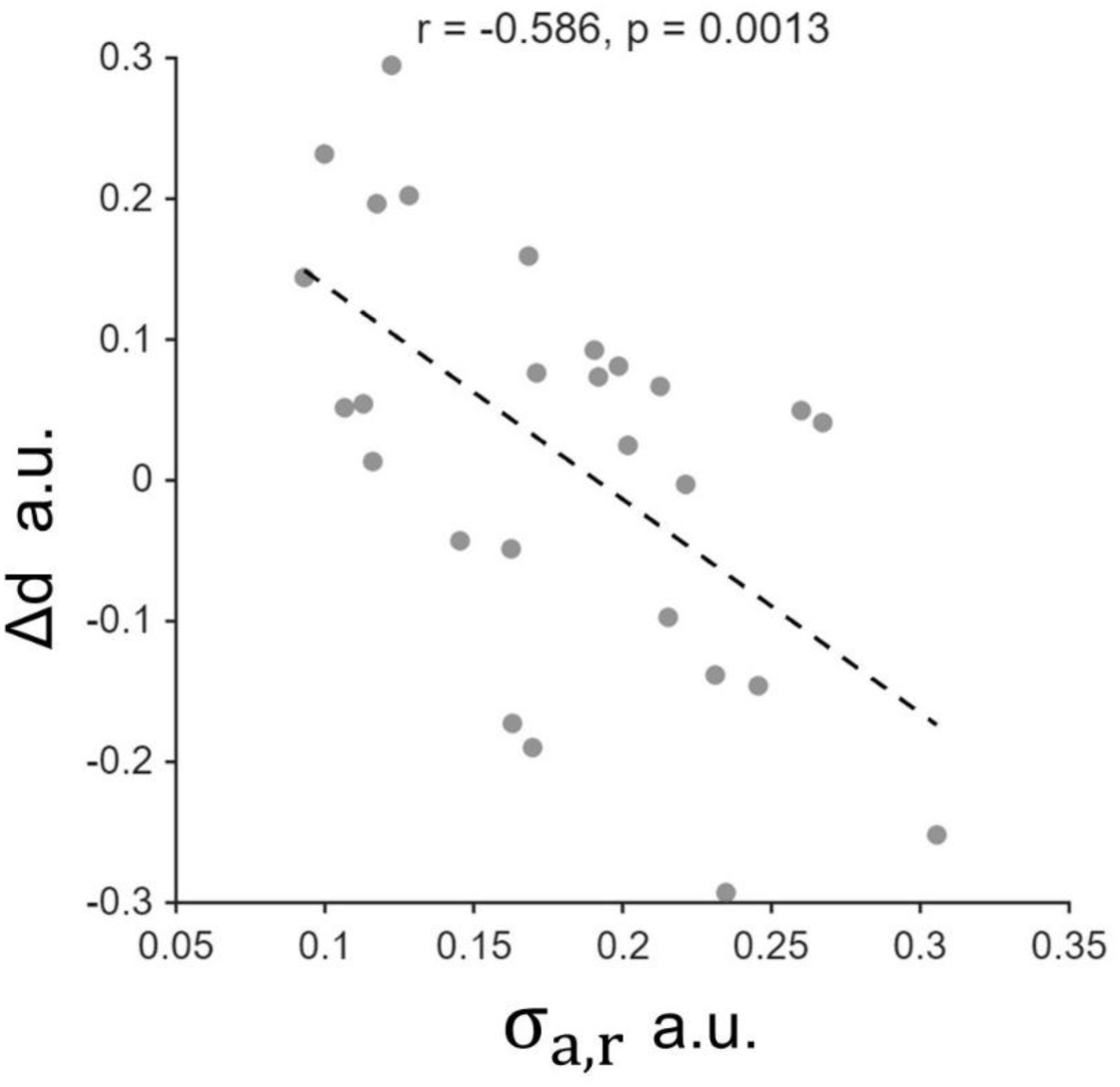
Relationship between individual variability and Δd. Scatter plot shows the correlation between the ROI-specific inter-subject variability parameter and the change in ISS group difference effect size upon ROI exclusion (Δd) across the 27 Stroop-ROIs.

**Supplementary Figure 4.**
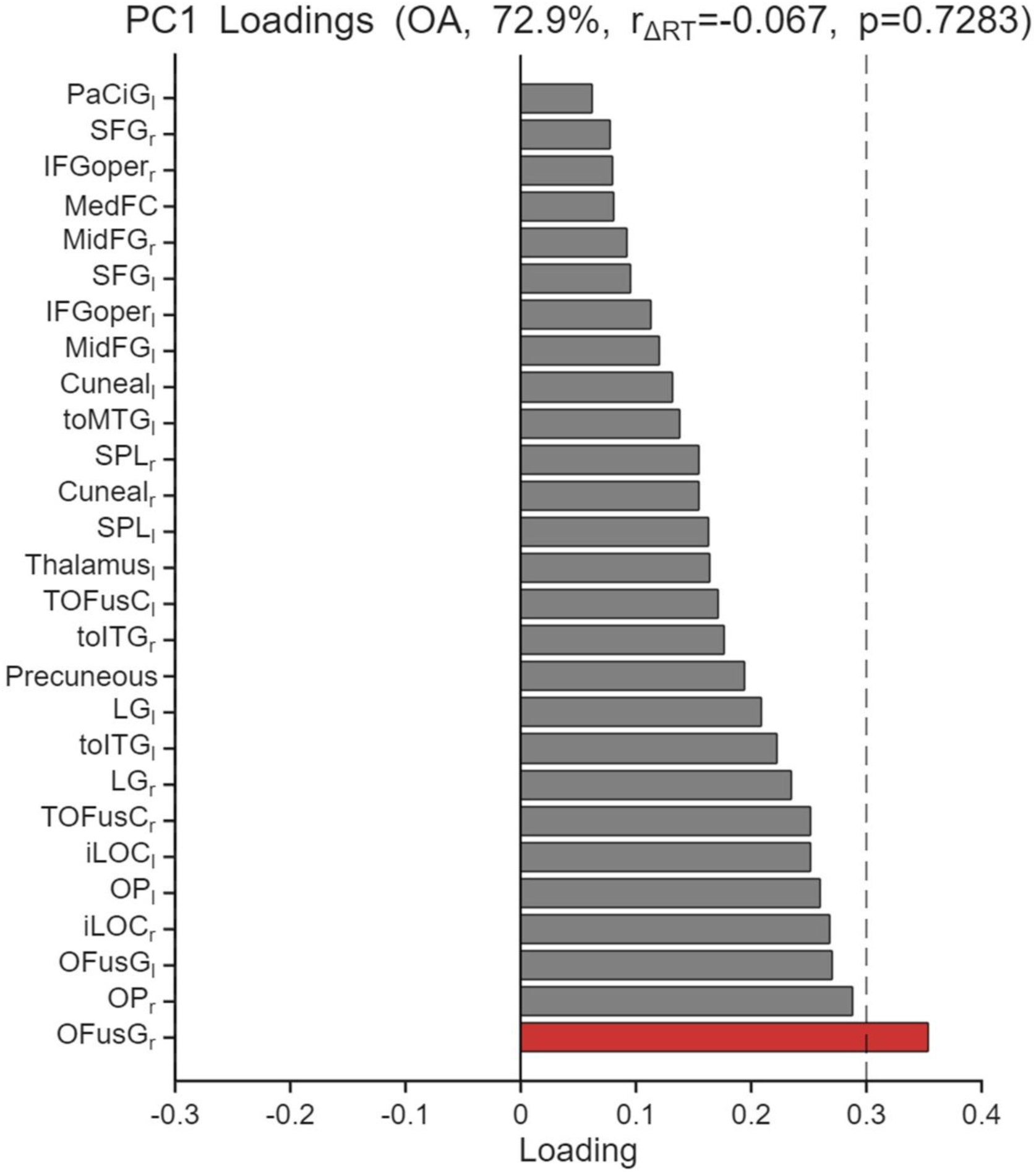
PC1 factor loadings across the 27 Stroop-ROIs. Factor loadings of PC1 derived from the subject-specific activation deviation matrix of the older group. Factor loadings with absolute values of 0.3 or greater are highlighted in red.

